# Divergent selection and primary gene flow shape incipient speciation of a riparian tree on Hawaii Island

**DOI:** 10.1101/698225

**Authors:** Jae Young Choi, Michael Purugganan, Elizabeth A. Stacy

## Abstract

A long-standing goal of evolutionary biology is to understand the mechanisms underlying the formation of species. Of particular interest is whether or not speciation can occur in the presence of gene flow and without a period of physical isolation. Here, we investigated this process within Hawaiian *Metrosideros*, a hyper-variable and highly dispersible woody species complex that dominates the Hawaiian Islands in continuous stands. Specifically, we investigated the origin of *Metrosideros polymorpha* var. *newellii* (newellii), a riparian ecotype endemic to Hawaii Island that is purportedly derived from the archipelago-wide *M. polymorpha* var. *glaberrima* (glaberrima). Disruptive selection across a sharp forest-riparian ecotone contributes to the isolation of these varieties and is a likely driver of newellii’s origin. We examined genome-wide variation of 42 trees from Hawaii Island and older islands. Results revealed a split between glaberrima and newellii within the past 0.3-1.2 million years. Admixture was extensive between lineages within Hawaii Island and between islands, but introgression from populations on older islands (*i.e.* secondary gene flow) did not appear to contribute to the emergence of newellii. In contrast, recurrent gene flow (*i.e.* primary gene flow) between glaberrima and newellii contributed to the formation of genomic islands of elevated absolute and relative divergence. These regions were enriched for genes with regulatory functions as well as for signals of positive selection, especially in newellii, consistent with divergent selection underlying their formation. In sum, our results support riparian newellii as a rare case of incipient ecological speciation with primary gene flow in trees.

**Author summary:** A long-standing question in evolution is whether or not new species can arise in the presence of gene flow, which is expected to inhibit the formation of reproductive isolating barriers. We investigated the genomics underlying the origin of a Hawaii Island-endemic riparian tree and purported case of incipient sympatric speciation due to disruptive selection across a sharp forest-riparian ecotone. We find extensive evidence of ongoing gene flow between the riparian tree and its closest relative along with local genomic regions resistant to admixture that likely formed through selection on genes for ecological adaptation and/or reproductive isolation. These results strongly suggest that where disruptive selection is strong, incipient speciation with gene flow is possible even in long-lived, highly dispersible trees.

## Introduction

A major aim of evolutionary biology is to understand the mechanisms underlying the formation of species. Of particular interest is the formation of species from within a panmictic population [1] or between populations that co-occur within geographic “cruising range” [2], as gene flow between diverging populations is expected to erode linkage between alleles associated with differential adaptation and reproductive isolation, thus reversing speciation [3]. As a result, the “speciation-with-gene-flow” model is highly controversial [4–6]. Moreover, when diverging populations are connected by ongoing (“primary”) gene flow without any period of physical isolation (i.e., allopatry), speciation is generally not expected [3,7–9]. In fact, recent analyses enabled by the availability of population-level genome sequence data have challenged the few purported cases of speciation with primary gene flow [10] by revealing historical periods of allopatry [11–13], thus indicating that gene flow had occurred through secondary contact [10, 14]. Regardless, despite the rarity of cases, speciation-with-primary-gene-flow (or sympatric speciation [14]) is an intriguing model of speciation for evolutionary biologists, one that requires the integration of both ecology and genetics for its inference [6].

Sympatric speciation occurs predominantly through strong disruptive selection [4, 15]. When incipient species inhabit different environments, divergent selection (*i.e.* selection acting in contrasting directions between populations or selection favoring phenotypes of opposite extremes [16, 17]) favors locally adaptive alleles that are beneficial in the alternative ecological niches. In the ecological speciation model [18], reproductive isolation evolves between populations as a consequence of ecologically based divergent selection (but see Coyne and Orr [19] for non-ecological mechanisms). For instance, when a pair of incipient species have adapted to contrasting environments, reproductive barriers can arise in the form of immigrant inviability, where selection acts against migrants in each environment [20]. In plants, immigrant inviability is a strong isolating mechanism [21–25], suggesting that ecological speciation may be a principal driver of incipient speciation by restricting gene flow between sympatric populations through a process analogous to reinforcement [20, 26]. The ecotypes that arise through divergent selection should represent the earliest stage of speciation [17]. These lineages are likely to be only partially reproductively isolated, which makes them ideal for studying the initial stages of barrier formation [27] before the onset of confounding genetic changes that accumulate once speciation is complete [28].

At the genomic level, under the ecological speciation-with-gene-flow model [27] gene flow will homogenize the genomic landscape and prevent the divergence of regions not directly under divergent selection or linked to such regions [16,29,30]. Intense interest has focused on identifying these genomic islands of divergence [31–33], since these regions should contain the genetic variants involved in speciation [32,34–41]. Any genomic region resistant to gene flow would form localized peaks of differentiation that can be readily detected through genome-wide scans of differentiation (*e.g.* F_ST_) [29, 42]. However, because differential selection rather than differential gene flow can also produce islands of divergence through genetic hitchhiking or background selection [43–45], absolute measures of differentiation such as D_xy_ have been proposed as an alternative for identifying islands of divergence [46, 47]. In plants, studies of the role of ecological divergence in the formation of reproductive barriers at the genic/genomic level are few and involve just a handful of “model” species [48]. Reconstructing the past demographic histories of the diverged populations and using the combined measures of F_ST_ and D_xy_ in multiple populations can aid our understanding of the genomic changes that accompany incipient speciation [49].

*Metrosideros* Gaud. (Myrtaceae) is a young (∼3.9 mya; [50]) woody species complex that dominates Hawaii’s native flora, comprising more than 20 vegetatively distinct [51, 52] but interfertile taxa that are non-randomly distributed across ecotones and environmental gradients on each of the main islands [51, 53]. Differential local adaptation across Hawaii’s heterogeneous landscape [54–61] and partial reproductive isolation [62, 63] have evolved among *Metrosideros* taxa despite their long life spans [64] and the high dispersibility of their seeds and pollen [54,65,66], two characters that are expected to impede diversification [67]. *Metrosideros* appears to be an example of incipient adaptive radiation with gene flow in trees [68].

*Metrosideros polymorpha* var. *newellii* (hereafter newellii) is an incipient species endemic to the waterways of the windward (wet) side of Hawaii Island. Newellii is a morphologically distinct, narrow-leaved ecotype (Fig 1A) that appears to derive from the archipelago-wide, broad-leaved *M. polymorpha* var. *glaberrima* (hereafter glaberrima; Fig 1A) [68] following a progenitor-derivative speciation model [69]. Riparian zones on Hawaii Island are narrow (typically the width of a single tree on either side of the waterway) and embedded within glaberrima-dominated wet forest. Along waterways, mating opportunities among adults of newellii and glaberrima are presumably random, consistent with a strict definition of sympatry [2,6,70–73], and apparent hybrids are occasionally found. Despite sympatry, glaberrima and newellii are surprisingly strongly isolated at SSR loci (F_ST_ = 0.094 (max = 0.16) relative to an average F_ST_ = 0.054 among all Hawaii-Island varieties except newellii [68]).

**Figure 1.**
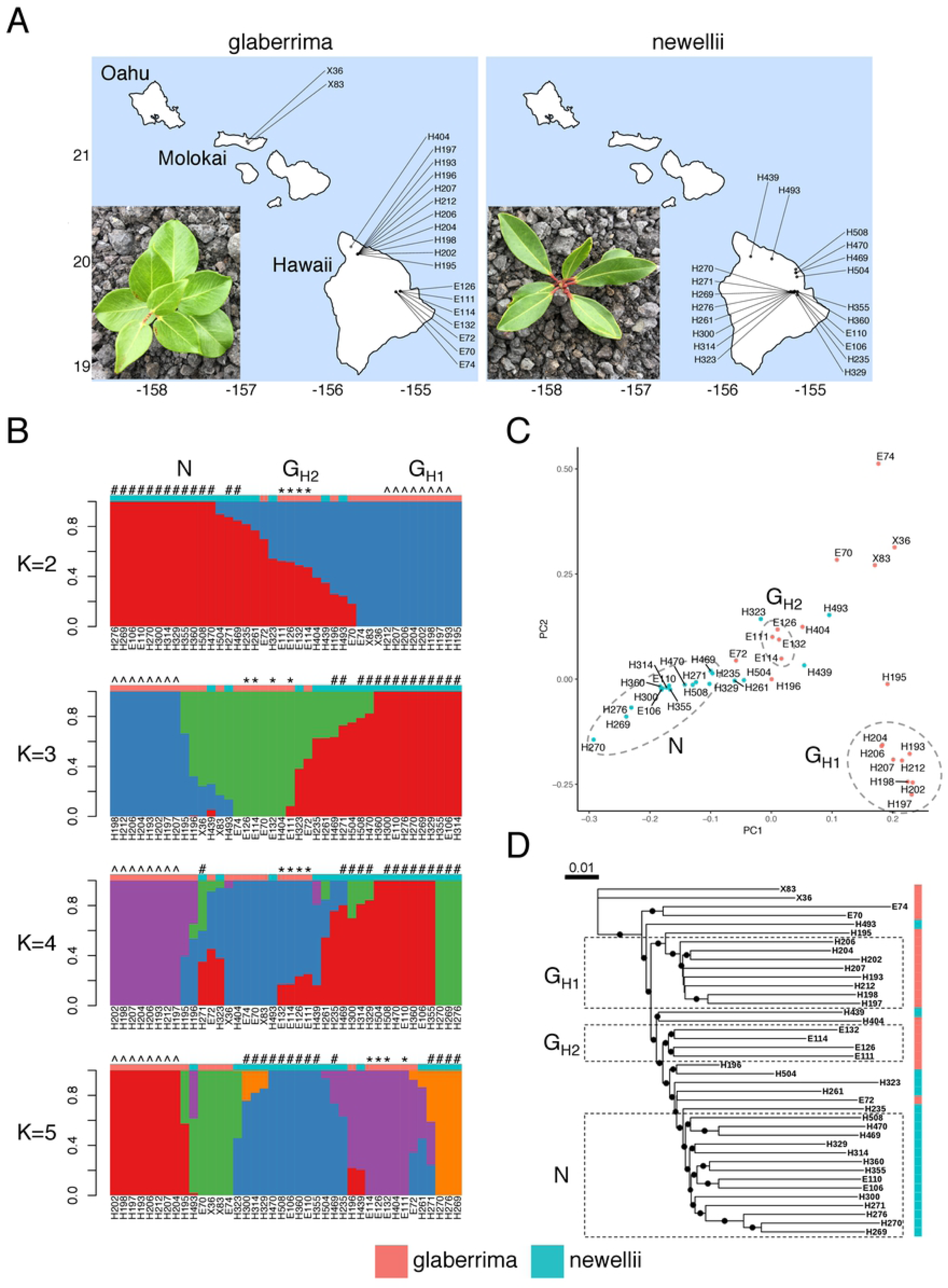
Genetic relationship between glaberrima and newellii, two varieties of *M. polymorpha*. G_H1_, G_H2_, and N are the genetic populations identified through this study. (A) Geographic locations of the 40 sampled trees of glaberrima and newellii. Inset shows the leaf morphology of glaberrima and newellii. (B) ADMIXTURE plot of K=2, 3, 4, and 5 for the 40 trees. Individuals from G_H1_, G_H2_, and N are indicated with ^, * and #, respectively. (C) PCA plot of the 40 trees. (D) Neighbor-joining tree of the 40 trees. Nodes with greater than 90% bootstrap support are indicated with black circles.

Riparian zones on Hawaii Island are harsh, and disruptive selection across a sharp ecotone results in partial reproductive isolation through significant reciprocal immigrant inviability [20] in adjacent forest and riparian environments [74]. Seedlings of newellii are better adapted than those of glaberrima to the strong water current and secondarily to the high-light stress of these environments [74]. Whether additional reproductive isolating barriers exist is not yet known. Regardless, the morphological and ecological divergence and strength of genetic differentiation observed between newellii and glaberrima equal or exceed those of other documented incipient species pairs or radiations (e.g [75–78]). Although newellii is endemic to Hawaii Island (0.5 mya [79]), an archipelago-wide analysis of SSR variation in Hawaiian *Metrosideros* suggested that it may have arisen from glaberrima on the prehistoric island of Maui nui, comprising modern-day Maui, Lanai, Molokai, and Kahoolawe (≤1.8 mya [79, 80]).

We investigated the demographic history of *M. polymorpha* during its colonization of Hawaii Island, the relationship between newellii and glaberrima, and the genomic regions that were involved in the origin of newellii. We found evidence of extensive admixture between close and divergent lineages, including evidence that incipient speciation between glaberrima and newellii is consistent with a speciation-with-primary-gene-flow model. Examination of the genomic islands of divergence that formed during incipient speciation revealed evidence of positive selection on genes associated with the regulation of gene expression, consistent with the origin of newellii through disruptive selection across a forest-riparian ecotone. We estimate that newellii emerged 0.3-1.2 mya.

## Results

Forty-two adults were chosen for genome sequencing, including 18 glaberrima and 20 newellii adults from Hawaii Island, two glaberrima adults from Molokai, and two outgroup species (*M. rugosa* and *M. tremuloides*) from Oahu (Fig 1A). Genome coverage was on average ∼15×, ranging from 1× to 43× per individual (S1 Table). For the majority of the samples more than 95% of the reads could be aligned to the reference *M. polymorpha* genome (S1 Table) [81]. Because the reference genome was still in a draft state with over 55,000 scaffolds, however, we restricted alignments to just scaffolds that were greater than 100,000 bps (totaling 285 Mbp of the 304-Mbp genome assembly) to avoid analyzing potential repetitive regions in unassembled contigs.

### Population structure of glaberrima and newellii on Hawaii Island

Because of the varying sequence depth among individuals, population relationships were examined using a complete probabilistic approach to account for the uncertainty in genotypes [82, 83]. Polymorphic sites were pruned randomly to minimize the effect of linkage between physically close sites.

Using the ADMIXTURE method [84, 85], ancestry proportions for each individual were estimated by varying the assumed number of ancestral populations (K) from 2 to 7 (S1 Fig). At K=2, individuals were largely divided into two groups representing the two varieties (Fig 1B). Principal components analysis (PCA) corroborated the K=2 ancestry result with the first component dividing the individuals by variety (Fig 1C). The phylogenetic analysis also revealed a near-complete division of individuals by variety (Fig 1D).

Genetic structure was observed within each variety as well (Fig 1). We used monophyly (Fig 1D) to identify three core groups (G_H1_, G_H2_, and N). G_H1_ comprised eight glaberrima trees from the Alakahi Bog on Kohala Volcano that formed a separate cluster in the principal component space (Fig 1C) and showed a unique ancestry in the ADMIXTURE analyses with higher K values (Fig 1B). G_H2_ comprised four closely spaced glaberrima trees situated alongside newellii at Kalohewahewa Stream. These four individuals, along with three others at the Wailuku_A site (E70, E72, E74) were glaberrima trees sampled immediately adjacent to newellii trees along waterways (Fig 1A). Phylogenetically, G_H2_ was sister to newellii (Fig 1D) and in principal component space was closer to newellii than was G_H1_ (Fig 1C). The ancestry proportions for G_H2_ were inconsistent across K values, but the results for K=2, 3 and 4 indicated that ancestry of G_H2_ included newellii and a glaberrima population that was not related to G_H1_.

Group N comprised 14 individuals of newellii that occur along the largest waterway on Hawaii Island, the Wailuku River. While individuals from the N group clustered together in principal component space (Fig 1C), there was evidence of substructure within the group. Trees from the most up-river population (Wailuku_A) were most diverged in the phylogenetic tree and formed their own cluster at higher K values (K=4 and 5; Fig 1B).

Among the individuals excluded from these three monophyletic groups were two glaberrima trees (E70 and E74) that were genetically similar to glaberrima from Molokai despite occurring alongside the most divergent newellii subpopulation (i.e., Wailuku_A; Fig 1). Hence, there may be further genetic structure with both varieties on Hawaii Island, but low sample sizes from these additional subgroups precluded further analysis. Lastly, two trees, H439 and H493, from northern waterways were phenotypically characterized as newellii, but were more closely related to the glaberrima group, and a single phenotypically defined glaberrima (E72) grouped within newellii (Fig 1).

### Admixture between glaberrima and newellii on Hawaii Island

We limited subsequent analyses to the G_H1_, G_H2_, and N populations, glaberrima from Molokai (individuals X83 and X36, hereafter referred as G_M_), and the two outgroup species, *M. rugosa*, and *M. tremuloides*. There were 8,436,114 variable positions across all samples, 7,566,381 variable positions within *M. polymorpha*, and 6,818,011 variable positions among the samples from Hawaii Island.

Consistent with the genotype likelihood-based tree (Fig 1D), the phylogenetic relationships from computationally-called genotypes indicated a tree topology of [(((G_H2_, N), G_H1_), G_M_), *M. rugosa*, *M. tremuloides*] (S2 Fig). Since admixture events occurring between branches are not detected in a simple bifurcating tree, we used methods that quantify, or test for evidence of, admixture occurring between lineages.

To characterize the relationships among G_H1_, G_H2_, and N on Hawaii Island we implemented a topology-weighting method [86]. Using one of G_M,_ *M. rugosa*, or *M. tremuloides* as the outgroup, we conducted sliding-window analyses and in each window quantified the taxon topology weight, which is a count of all of the unique sub-trees within which a single individual of each taxon is represented. Initially, results showed that regardless of the outgroup used, no single topology dominated the weighting (Fig 2A), suggesting that many variants are shared among G_H1_, G_H2_, and N. The choice of outgroup, however, did affect the majority topology. Specifically, with G_M_ or *M. rugosa* as the outgroup, the topology that grouped N and G_H2_ as sisters had the highest weighting (Fig 2A dotted box). In contrast, with *M. tremuloides* as the outgroup the topology that grouped N and G_H2_ as sisters had the lowest weighting, consistent with admixture between *M. tremuloides* and either G_H2_ or N.

**Figure 2.**
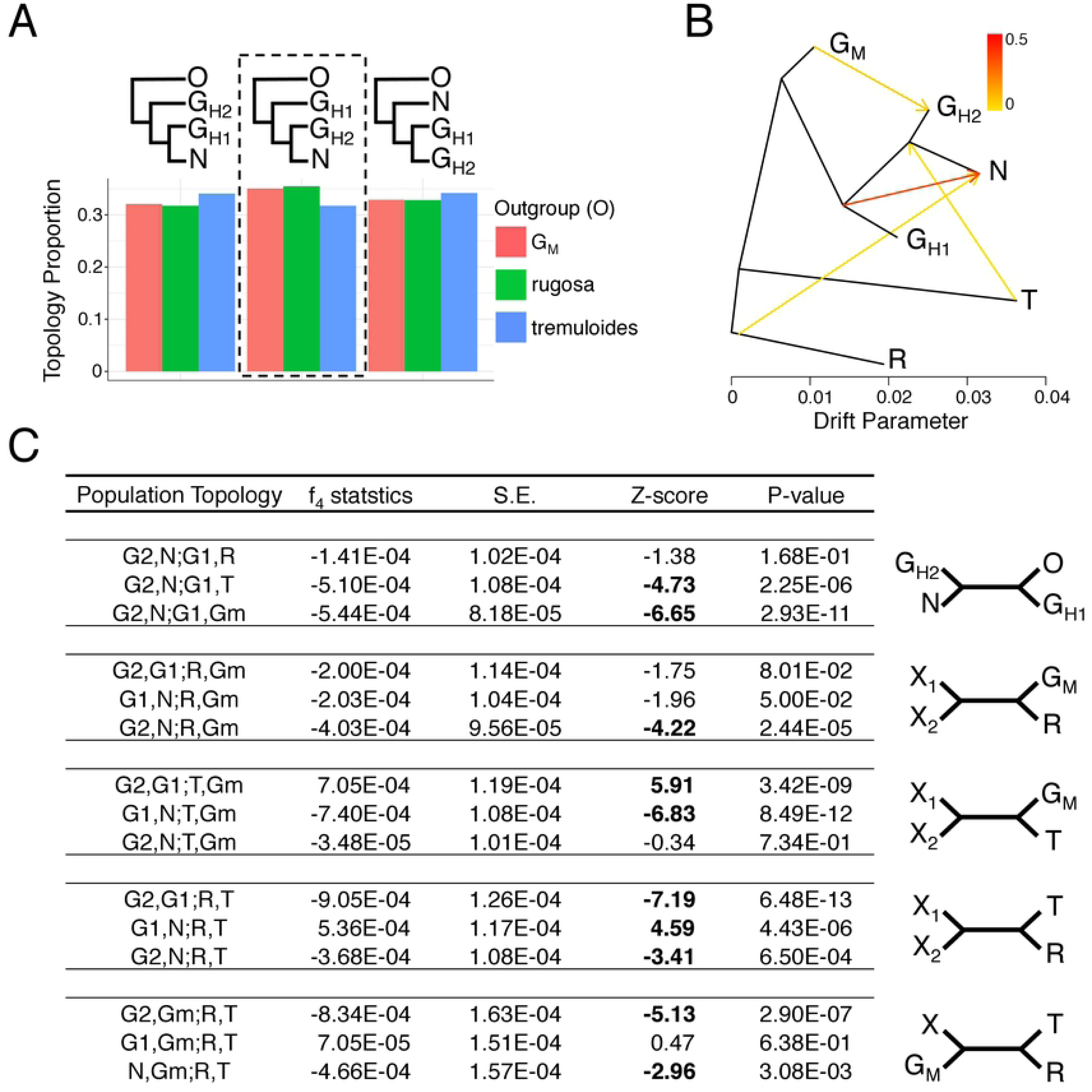
Relationships among the three focal populations of *M. polymorpha* on Hawaii Island. (A) Topology weight for a three-population topology under three different outgroups. Dotted box represents the topological relationship obtained from genome-wide data. (B) Treemix graph of the three focal populations and three outgroups assuming 4 migration edges. (C) Results of f_4_ tests of admixture in the m=4 treemix graph.

Admixture among the three Hawaii Island populations and each of the outgroup populations was examined using treemix [87] by fitting migration edges on a bifurcating tree (Fig 2B). The initial treemix tree with zero migration recapitulated the population relationships recovered using the topology-weighting method, while the treemix graphs with increasing numbers of migration edges showed different relationships (S3 Fig). Briefly, the first migration edge was fitted between *M. tremuloides* and the common ancestor of G_H2_ and N, consistent with the topology-weighting results. At two migration edges, there was admixture between G_H1_ and N, whereas in higher migration models the common ancestor of all three Hawaii Island populations admixed into N. At three migration edges, there was admixture between G_M_ and G_H2_, while at four migration edges, an unsampled *M. rugosa-*like population was admixed with N. At five migration edges, there was additional gene flow originating from the common ancestor of the Hawaii Island populations into G_H2_. This suggests that five migration edges may over-fit the data or that Hawaii Island *M. polymorpha* may have originated as a hybrid swarm [88–90].

The treemix results suggested that N and G_H2_, the two sister groups from Hawaii Island, had extensive evidence of admixture with other populations from both Hawaii Island and one or more older islands. To specifically test for evidence of introgression we implemented the four-population (f_4_) test [91]. All significant f_4_ test results were consistent with the treemix result with four migration edges (Fig 2C) and revealed two major admixture patterns. (1) The significant f4 test on the topology [(G_H2_, N);(G_H1_, T or G_M_)] was consistent with evidence of admixture between N and G_H1_, and admixture between G_H2_ and G_M_. The non-significance of the topology [(G_H2_, N);(G_H1_, R)] is likely due to N but not G_H2_ sharing alleles with both G_H1_ and *M. rugosa*. (2) The significant f_4_ test on topology [(G_H2_ or N, G_H1_);(R, T)] was consistent with *M. tremuloides* admixing with either G_H2_ and N, or their common ancestor. This also indicated that among the three Hawaii Island groups, G_H1_ was the least affected by admixture from lineages on older islands.

### Demographic history of Hawaii Island glaberrima and newellii

Initially, we focused on the demographic histories of G_H1_ and N to infer the recent splitting event that led to the formation of newellii. Levels of polymorphism were similar between the two groups (median θ_π_ = 0.0041 and 0.0043 for G_H1_ and N, respectively), while linkage disequilibrium decreased rapidly to low levels (r^2^ < 0.2) within 1 kbp for both populations (S4 Fig). Median F_ST_ between G_H1_ and N was 0.043 (S5 Fig).

The demographic histories of G_H1_ and N were estimated using the diffusion approximation method of dadi [92]. Twenty demographic models (S6 Fig) [93] were fit onto the observed two-dimensional joint-site-frequency spectrum (2D-SFS) between G_H1_ and N (see S2 Table for all models with their estimated parameters and log-likelihood). The top three best-fitting models for the 2D-SFS all involved an ongoing or secondary contact-based gene flow between G_H1_ and N. In addition, the demographic model with recent or ongoing gene flow was chosen as the best-fitting model regardless of whether the underlying 2D-SFS was estimated using genotype likelihood or genotype calls (see S3 Table for dadi results using genotype calls). The top three best-fitting models were then further optimized to fit the best-fitting demographic model (S4 Table). In the end the best-fitting model was one that modeled an ongoing asymmetric migration between G_H1_ and N (ΔAIC=18.8 and see S7 Fig for model fit). The model estimated a higher level of gene flow from G_H1_ into N, while the effective population size of G_H1_ was slightly higher than that for N (Fig 3A).

**Figure 3.**
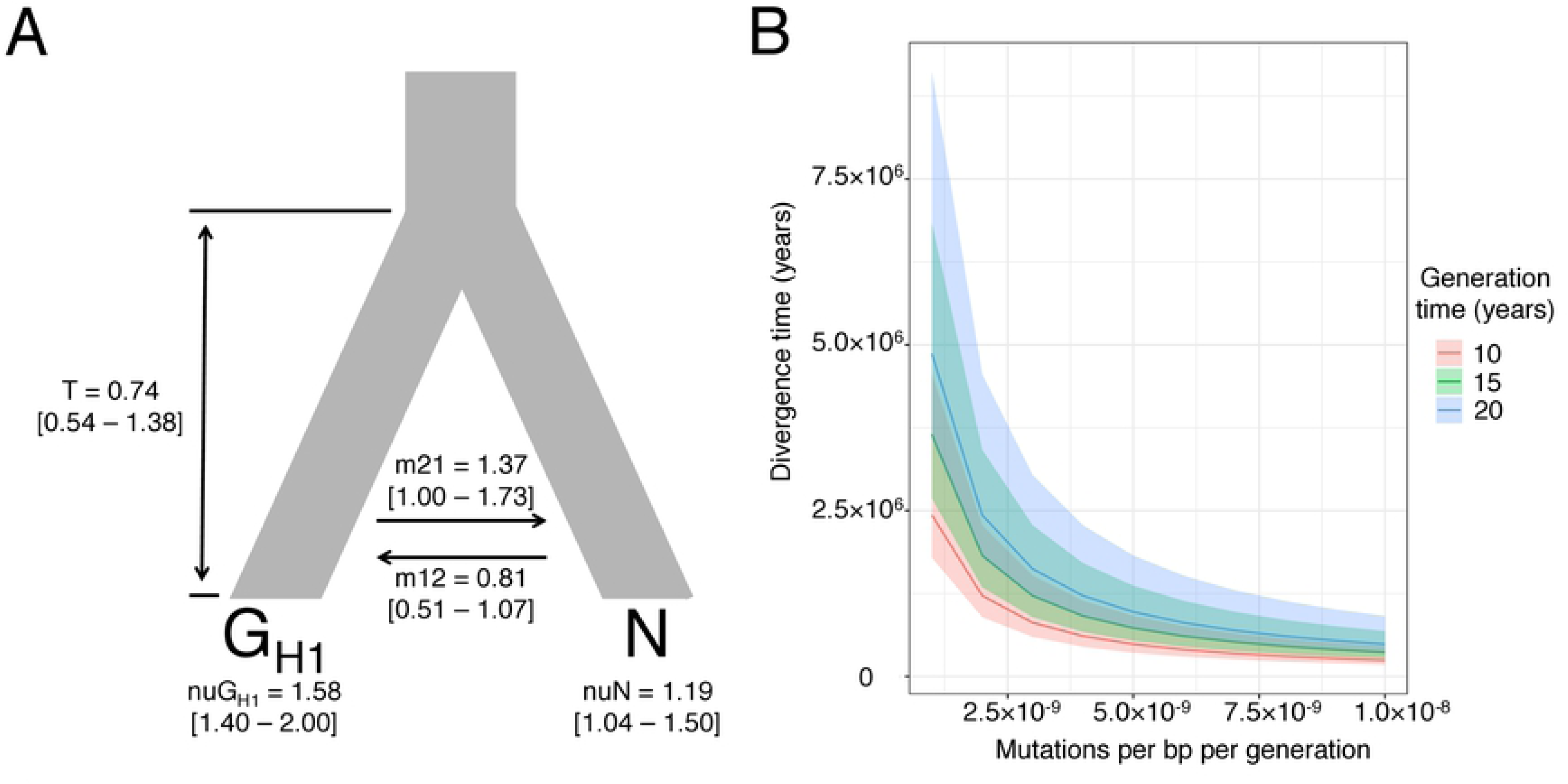
Estimation of demographic parameters for newellii and G_H1_ (the glaberrima closely related and purported progenitor to newellii) on Hawaii Island. (A) dadi parameter estimates for the best-fitting model. Effective population sizes of current populations are scaled by the effective population size of the ancestral population (N_anc_) that is arbitrarily set to 1. Time is reported in unscaled units of T where T×2N_anc_=τ (divergence time in generations). Migration rates are reported in units of m_12_ where m_12_/2N_anc_= M_12_ (proportion of individuals in each generation that are new migrants from population 2 to population 1). Bootstrap parameter estimates are shown in square brackets. nuG_H1_: effective population size of G_H1_; nuN: effective population size of N; m12: migration rate from N to G_H1_; m21: migration rate from G_H1_ to N; and T: unscaled divergence time between G_H1_ and N. (B) Divergence time (years) between G_H1_ and N under various mutation rates and generation times. Bootstrap confidence intervals are shown in shaded areas.

We then tested if different proportions of the ancestral population contributed to G_H1_ and N, specifically testing whether N was derived from a smaller ancestral proportion than G_H1_ (*i.e.* an island model [94]). Seven different scenarios (S8 Fig) were tested, and all models indicated almost equal proportions of the ancestral population contributed to G_H1_ and N (S5 Table). Further, the best-fitting scenario was still less fit than the model of a simple split with continuous asymmetric gene flow (ΔAIC=5.42 and S6 Table), suggesting no evidence of a vicariance-like event between G_H1_ and N.

We also used dadi to estimate the demography between G_H2_ and N by testing the same 20 demographic models (see S6 Fig for models and S7 Table for initial optimization results). The best-fitting model was a two-epoch model of asymmetric migration (S8 Table), and the resulting unscaled divergence time (T) almost overlapped the bootstrap confidence interval of the divergence time for G_H1_ and N.

The T between G_H1_ and N can be converted to years (*i.e.* scaled divergence time (τ)) with a mutation rate and generation time, both of which are unknown for *Metrosideros.* Hence, we estimated τ assuming a range of mutation rates (*Arabidopsis* mutation rate = 7×10^-9^ [95], *Prunus* mutation rate = 9.5×10^-9^ [96]) and generation times reasonable for these trees (Fig 3B). τ varied the most at lower mutation rates, but within previously known plant mutation rates (>7×10^-9^ mutations per bp per generation) all estimated divergence times were within 1.25 million years regardless of generation time.

### Genomic islands of divergence between G_H1_ and N

The genomic landscape of differentiation between G_H1_ and N was estimated through local genomic windows of F_ST_ values. F_ST_ was Z-transformed (zF_ST_), and genomic regions with zF_ST_ > 4 were considered significant outliers [35] (Fig 4A). Genomic islands with increased levels of relative divergence (F_ST_) were identified in 237 out of 27,508 windows (merged into 147 non-overlapping windows) for a total of 2.37 Mbp. Further, compared to the genomic background, these islands had significantly elevated levels of absolute divergence (D_xy_) (Fig 4B; Mann Whitney U [MWU] p-value = <1e-10). In addition, when the regions of genomic islands identified between G_H1_ and N were examined between G_H1_ and G_H2_, D_xy_ values were also significantly elevated compared to the genomic background, indicating that islands of divergence are shared between G_H2_ and N. Between G_H2_ and N, however, D_xy_ was significantly lower in the genomic islands identified between G_H1_ and N (S5 Table).

**Figure 4.**
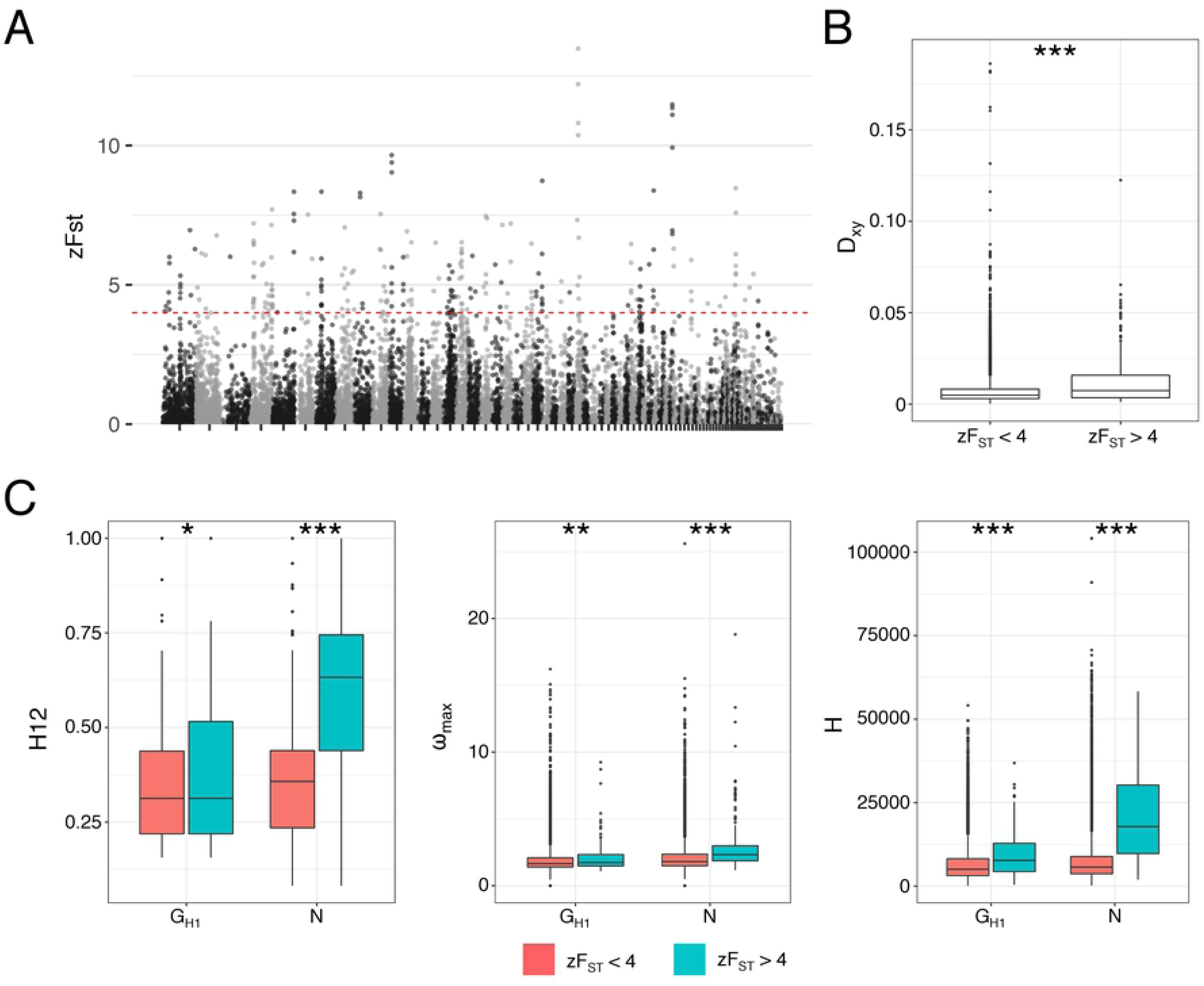
Patterns of genome-wide divergence between G_H1_ and N. (A) Genome-wide zFst values calculated from 10-kbp windows. Scaffolds were ordered from longest to shortest. (B) Absolute measures of divergence (D_xy_) between genomic islands of divergence (zF_ST_ > 4) and the genomic background (zF_ST_ < 4). (C) Selective sweep test statistics for genomic islands of divergence and the genomic background. Significant differences are indicated with * < 0.05, ** < 0.01, and *** < 0.001.

Levels of polymorphism were significantly lower in genomic islands relative to background in both G_H1_ and N (S9 Fig; MWU p-value = <1e-15), suggesting that the increased F_ST_ and D_xy_ were partly caused by those regions being frequent targets of selection. We measured haplotype homozygosity (H12) [97] and haplotype length (H) [98], and performed an LD-based test of selection (ω_max_) [99], and found all selective sweep statistics were significantly elevated in genomic islands (Fig 4C; MWU p-value = <1e-10) for both G_H1_ and N. In addition, N seemed to have more genomic islands with evidence of a selective sweep, shown by the overall higher levels of selective sweep statistics for N relative to G_H1_. We note that recurrent sweeps from a common ancestor are expected to lead to a decrease in D_xy_ [46]. Our observations of increased D_xy_ and evidence of selection, in contrast, suggest that islands of divergence were shaped by positive selection on divergent ancestral haplotypes predating the split between G_H1_ and N [100].

We then examined if the secondary contact between G_H1_ and/or N and outgroup populations on older Hawaiian Islands (G_M_, *M. rugosa*, and *M. tremuloides*) had contributed to the formation of the genomic islands of divergence. If secondary gene flow was recurrent, the genomic islands of divergence identified between G_H1_ and N would also have elevated D_xy_ levels between G_H1_ or N and the relevant outgroup. However, no such effect was observed (S9 Table), suggesting that sufficient levels of genome-wide divergence had eroded the genomic islands in allopatric comparisons [30]. We then further examined the N group because of its extensive evidence of admixture with outgroup species, in particular *M. tremuloides*, and its incipient speciation status. We tested whether genomic islands had origins relating to *M. tremuloides* by estimating localized windows of f_4_ statistics for the topology [G_H1_,N;G_M_,tremuloides]. Because G_H1_ was least affected by admixture with any of the outgroups (Fig 2), an excess of positive f_4_ statistics across local windows would indicate increased admixture between N and *M. tremuloides* in the region. There was no significant difference in f_4_ statistics between the genomic islands of divergence and the genomic background. In addition, we calculated the genic proportion of each window to examine if the admixture had affected coding sequences more than non-coding regions, suggesting a role for selection maintaining the introgression. No significant correlation was seen between the proportion of gene sequence per window and either positive or negative f_4_ statistics (S10 Fig). These results, in the end, suggested that secondary contact with allopatric lineages did not contribute to the incipient speciation of N.

Gene ontology (GO)-enrichment analysis was conducted for the 341 genes (S10 Table) that overlapped with genomic islands of divergence. GO could be assigned to 147 genes, which were significantly enriched (hypergeometric test p-values < 0.05) for functions relating to two major categories: 1) DNA binding and transcriptional regulation (GO:0003700, GO:0140110, and GO:0003677) and 2) acetyltransferase activity (GO:0016747 and GO:0016407).

## Discussion

Newellii on Hawaii Island appears to be a case of early-stage incipient speciation in a highly dispersible and long-lived tree. Based on our evidence, we argue that divergence between newellii and glaberrima is consistent with a speciation-with-primary-gene-flow model, where barriers to gene flow are developing through disruptive selection. The incipient status of newellii notwithstanding, our results place this riparian tree in the small but growing number of cases of speciation occurring despite the homogenizing effects of gene flow. This pattern is consistent with the observation of differential adaptation of newellii to the mechanical stress of rushing water and high light. [74] and the greater shade tolerance of wet-forest-dominant glaberrima [55] resulting from disruptive selection across the forest-riparian ecotone.

Evidence of admixture was extensive among the sampled populations, occurring within and between different island lineages, and may be common throughout the species complex [101]. Our results are in line with the recent view of sympatric speciation in the genomic era, in which gene flow is almost ubiquitous between recently diverged sister species [14, 102]. Clarifying the relative roles of primary versus secondary gene flow during speciation, however, is critical for understanding the evolutionary processes underlying the divergence between sympatric sister lineages [14]. For instance, our genomic analyses suggested a close genetic relationship between G_H2_ and N, yet one that did not result from divergence in sympatry. N and G_H2_ shared islands of divergence, but these regions were not elevated in D_xy_ between N and G_H2_. This result suggests that the shared islands of divergence between N and G_H2_ arose through recent shared selective sweeps [46], excluding the possibility of primary gene flow (and subsequent divergent selection) underlying their formation. Given the close proximity of N and G_H2_, the appearance of locally adaptive alleles from N in G_H2_ may result from introgression between varieties where the forest-riparian ecotone is more diffuse and disruptive selection is less intense [74]. The recent admixture between G_H2_ and G_M_ is also consistent with the formation of G_H2_ through secondary contact between a G_M_-like lineage and N. Our results also indicated gene flow between N and an allopatric lineage (*M. tremuloides* on Oahu). The genomic regions resulting from this secondary contact, however, were not enriched in the islands of divergence between G_H1_ and N. This result suggests that primary gene flow between G_H1_ and N, but not secondary gene flow between N and *M. tremuloides*, contributed to the formation of genomic islands of divergence. While this secondary gene flow may have been neutral during speciation, it is possible that some of the introgressed regions were utilized during local adaptation unrelated to incipient speciation [34,103–107].

The divergence time between glaberrima and newellii was estimated to be between 347-695 kya (assuming a mutation rate of 7×10^-9^ bp per site per generation and generation time of between 10 to 20 years). This time frame coincides roughly with the geological age of Hawaii Island, which formed ∼500 kya [79], and is consistent with an in-situ origin of newellii. Our more conservative estimates of divergence time, however, extend back to ∼1.2 mya and suggest that divergence between glaberrima and newellii may have begun on the next youngest island in the chain, Maui Nui, before the colonization of Hawaii Island. While the site of origin of newellii remains unresolved, our results suggest that Hawaii-Island glaberrima-newellii may have derived from a hybrid swarm-like population (Fig 2). Harsh environments such as newly formed volcanic islands may promote colonization by hybrid progenies [108] with transgressive traits that are not seen in either parent [109, 110]. A denser sampling of *M. polymorpha* on the islands of Maui Nui may distinguish whether the Hawaii Island glaberrima-newellii complex has a deeper ancestry (*i.e.* hybridization of multiple progenitors) or a recent ancestry (*i.e.* deriving from a single ancestral background).

Genome-wide F_ST_ levels suggested that gene flow has largely homogenized the genomes of newellii and glaberrima, but we discovered regions resistant to gene flow based on elevated absolute and relative divergence, and these genomics islands of divergence were enriched for evidence of positive selection in both varieties. Given the contrasting ecological niches of newellii and glaberrima [68, 74], it seems likely that these islands of divergence are associated with ecological divergence between the varieties [111, 112]. Our results further suggested that selection may have been stronger or more frequent in the island-endemic newellii, consistent with isolation by adaptation to Hawaii Island’s harsh riparian environment. These islands of divergence were enriched for genes with functions related to transcription factor activity, suggesting that divergence in the gene expression landscape underlies the emergence of this riparian specialist. Indeed, many “speciation genes” are known to have functions relating to DNA binding and transcriptional regulation [113, 114], and global gene expression patterns of hybrids from reproductively isolated species are often mis-regulated [115]. Future work is needed to understand the changes in genetic architecture that coincide with incipient speciation of newellii.

## Materials and Methods

Sequencing data generated from this study were deposited under NCBI project ID PRJNA534153. Codes used in the analysis can be found from github (https://github.com/cjy8709/Hawaii_Metrosideros_polymorpha) and data generated from this study can be found in dryad.

### Sample genome sequencing

Samples analyzed within this study can be found in S1 Table. DNA was extracted from leaf buds and 2×100bp paired-end libraries were constructed using Nextera library kit. Libraries were sequenced on an Illumina HiSeq 2500 system.

Sequencing reads were aligned to the reference *Metrosideros polymorpha* var. *glaberrima* genome sequence from Izuno et al. [101]. FASTQ reads were aligned using the program bwa-mem version 0.7.16a-r1181 [116], and duplicate reads from PCR during the library preparation step were removed using picard version 2.9.0 (http://broadinstitute.github.io/picard/). Alignment statistics were reported using GATK version 3.8–0 (https://software.broadinstitute.org/gatk/) and samtools version 1.9.

### SNP calling and genotyping

Genotype calls were executed using GATK. Alignment BAM files were used by GATK HaplotypeCaller engine with the option ‘-ERC GVCF’ to output variants as the genomic variant call format (gVCF). The gVCFs of each sample were merged together to conduct a multi-sample joint genotyping procedure using the GATK GenotypeGVCFs engine. Variants were divided into SNPs and INDELs using GATK SelectVariants engine, and variant filtration were conducted using GATK VariantFiltration enigne with GATK bestpractice hard filter guidelines. All SNPs located within 5bps of an INDEL variant and any SNPs for which less than 80% of the samples were genotyped were removed.

### Population relationship analysis

Initially, genetic relationships between and within the individuals were examined using genotype likelihoods under a complete probabilistic framework using ANGSD version 0.929 [83] and ngsTools [82]. For all analyses we required a minimum base and mapping quality score of 30, while requiring genotypes from a minimum of 75% of individuals per SNP site, a minimum population read coverage of one-third the average population read depth, and a maximum population read coverage of three times the average population read depth. Because no adequate outgroup species genome sequence is available for *M. polymorpha* we implemented a folded site frequency spectrum. To minimize the effect of linkage on the population relation inferences we randomly pruned the polymorphic sites. With a sliding window of size 10,000 bp, a polymorphic site was randomly chosen, while requiring a minimum distance of 5,000 bps between adjacent SNP sites.

Ancestry proportions (K) were estimated with NGSadmix [85] using genotype likelihoods calculated from ANGSD. K was estimated from 2 to 7, and for each K the analysis was repeated 100 times to choose the run with the highest log-likelihood. Genotype posterior probabilities (GPP) were calculated with ANGSD for use in principal components analysis (PCA) and phylogeny reconstruction; GPP were translated to genetic distances between individuals using ngsCovar [82] for PCA and NGSdist [117] for genetic distances. Using the genetic distances, FastME ver. 2.1.5 [118] was used to reconstruct a neighbor-joining tree, which was visualized using iTOL version 3.4.3 (http://itol.embl.de/). Bootstrap resampling of the GPP was conducted using NGSdist generating a 1,000-bootstrap resampled dataset. Each resampled data set was used to estimate genetic distances, which were then used to estimate the bootstrap confidence level for the phylogenetic tree.

Genotype calls from Hawaii Island individuals plus the outgroups, *M. rugosa* and *M. tremuloides*, from Oahu were used to generate a neighbor-joining tree (S2 Fig). Genetic distance between two individuals (X and Y) was estimated using the Kronecker delta function-based equation [119]:

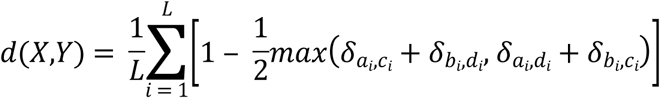

where *L* is the number of sites, *a_i_*and *b_i_* are the two allele copies in sample X, *c_i_*and *d_i_* are the two allele copies in sample Y, and *δ_jk_*is the Kronecker delta function (equals one if allele *j* is identical to allele *k* and 0 otherwise). From the genetic distance matrix, FastME was used to build the neighbor-joining tree.

### Tests for admixture

The local genomic topological relationship was examined using the topological weighting procedure implemented in twisst [86] (https://github.com/simonhmartin/twisst). Initially, local phylogenetic trees were estimated in sliding windows with a size of 100 polymorphic sites using RAxML with the GTRCAT substitution model. The script raxml_sliding_windows.py from the genomics_general package by Simon Martin (https://github.com/simonhmartin/genomics_general/tree/master/phylo) was used. We then used the ‘complete’ option of twisst to calculate the exact weighting of each local window by considering all subtrees.

Treemix version 1.13 [87] was used to fit migration edges on a bifurcating tree. We thinned the polymorphism data by randomly sampling a SNP every 1 kbp using plink version 2.0 to minimize the effects of LD between SNPs. The four-population test [91] was conducted using the fourpop program from the Treemix package.

### Demography modeling with dadi

Demography models were tested using the diffusion approximation method of dadi [92]. A visual representation of the 27 demographic models we tested can be found in S6 Fig and S8 Fig. Because we lacked an appropriate outgroup, folded 2-dimensional site frequency spectra (2D-SFS) were analyzed. 2D-SFS were generated using two different approaches: 1) using ANGSD to calculate the likelihood of the sample allele frequency followed by realSFS from the ANGSD package to calculate the expected number of sites within a given allele frequency based on that likelihood, and 2) using the genotype calls to calculate the allele frequency. With ANGSD the random polymorphic sites that were chosen in the “Population relationship analysis” were used to calculate the site allele frequency. With the genotype call dataset we randomly sampled a SNP every 10 kbp using plink and used this thinned variant call dataset. The script easySFS.py (https://github.com/isaacovercast/easySFS) was then used to generate the site frequency spectrum and choose the best sample size to project down and maximize the number of segregating sites to be analyzed.

Optimization of the model parameters was performed through 4 rounds of randomly perturbing the parameter values using the Nelder-Mead method. In the first round, random starting parameters were three-fold perturbed for a total of 10 replicates with a maximum of 3 iterations. Each optimized parameter was then used to simulate the 2D-SFS, and a multinomial approach was used to compare and estimate the log-likelihood of the observed 2D-SFS. Best-scoring likelihood was used in the second round for 20 replicates with parameters perturbed two fold and maximum iterations of 5. The best likelihood model of the previous round was used in a third round for 30 replicates with parameters perturbed two-fold and maximum iterations of 10. In the final round the best likelihood model of the third round was used for 40 replicates, with parameters perturbed one-fold, and maximum iterations of 15. Parameters from the round with the highest likelihood were selected to represent each demographic model. Akaike Information Criteria (AIC) values were used to compare demography models. The three best-fitting models were further analyzed with increased replicates and maximum iteration values. Specifically, with increasing rounds the replicates were 20, 40, 50, and 70; with maximum iterations of 10, 10, 25, and 50; and 3, 2, 2, and 1 fold parameter perturbations. The demography model with the lowest AIC was chosen as the best-fitting model.

Because ANGSD-based and genotype call-based site frequency spectra gave largely concordant results, all further analysis used only the ANGSD-based site frequency spectrum. Confidence of the best-fitting demography model parameter values was obtained through bootstrapping the site frequency spectrum and re-estimating the demography parameter values using the bootstrap data. The bootstrap site frequency spectrum was obtained using the realSFS program and randomly selected polymorphic sites by chromosome.

Divergence time between G_H1_ and N was a main parameter of interest, and the unscaled divergence times, *T*, were converted into years using the following equation:

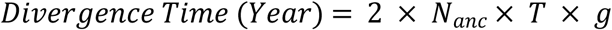

where N_anc_ is the ancestral population size and g is the generation time in years. *N_anc_* is unknown but can be inferred from the effective mutation rate θ (4N_anc_μL, where μ is the mutation rate and *L* is the total length of the sequenced region that was examined to analyze the SNPs), which was calculated by dadi in the optimal demographic model.

Here, the previous equation can be rewritten as:

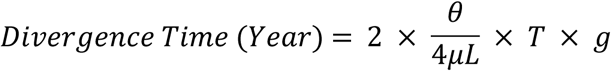

For *L*, ANGSD had analyzed 280,279,511 positions to detect polymorphic sites in 2,261,660 positions. But because we pruned the sites to 28,742 positions *L* was calculated as 280,279,511 × (28,742/2,261,660). Because *μ* and *g* for *M. polymorpha* are unknown, various biologically plausible values were used to estimate the divergence time.

### Population genetic analysis

The gVCFs from the SNP-calling step were used again for the GATK GenotypeGVCFs engine but to call genotypes for all sites including the nonvariant positions (option ‘--includeNonVariantSites’). This was done in order to correctly infer the number of variable and invariant sites for downstream calculation of window-based population genetics statistics. The genomics_general package was used to calculate θ, D_xy_, and F_ST_ in 10-kbp sliding windows and 5-kbp increments. Within each window sites were required to have a minimum quality score of 30 and a minimum depth of 5×. Only windows that had a minimum of 3 kbp of sites that passed the quality filter were further analyzed. For the calculation of sliding windows of f_4_ statistics, we used the script from Richards and Martin [106] and analyzed 10-kbp windows that had a minimum of 50 polymorphic sites.

Using the genotype calls, plink was used to calculate linkage disequilibrium (LD). LD was measured in squared correlations (r^2^) between polymorphic sites that were less than 3 kbp apart.

Evidence of selective sweeps was detected using haplotype-based or LD-based methods [97–99]. These approaches were taken because with no appropriate outgroup sequence it was not possible to polarize the variants and infer the high-frequency alleles, which are necessary for many allele frequency-based methods of detecting selective sweeps. The focal populations G_H1_ and N were phased and imputed using the program beagle ver. 5.0 [120]. Sweeps were detected using the haplotype homozygosity (H12), haplotype length (H), and LD-based tests of selection (ω_max_).

For the calculation of ω_max_ statistics, the total number of grids varied across scaffolds [121]. Grids were set so that ω_max_ would be calculated every 5,000 bp, and for each grid the minimum and maximum window size was set at 5,000 bp and 100,000 bp respectively. The H12 statistic was calculated by setting the analysis window size to 100 SNPs and sliding the analysis window by 10 SNPs. The H statistics were estimated for every SNP position. For any given window, statistics were averaged across all haplotype-based tests of selection to represent the selective sweep value for that window.

### Gene ontology enrichment

Coding sequences of each gene model were assigned a computationally predicted function and gene ontology using the eggnog pipeline [122]. An ontology could be assigned to 12,448 genes out of the total 37,956 gene models. We required an ontology to have more than 1 gene group member for further consideration. Gene ontology enrichment was tested through a hypergeometric test.

### Multiple testing corrections

All statistical tests with a p-value (Mann Whitney U test, Pearson’s correlation, f_4_ test, jack-knife bootstrap p-value, and hypergeometric test of gene ontology enrichment) were pooled together and corrected for multiple hypotheses testing using the Benjamini and Hochberg [123] correction method.

## Acknowledgements

We thank the Hawaii Department of Forestry and Wildlife for permission to collect leaf samples from state forests and J. Johansen for assistance with sample collection. We are grateful also to T. Sakishima, A. Veillet, and the Core Genetics Facility at the University of Hawaii Hilo for assistance with DNA isolation. We are also grateful to S. Ferrand at New York University Abu Dhabi (NYUAD) for assistance with the library preparation, and M. Gros-Balthazard at NYUAD for support with the genomic data. We thank E. Richards at University of North Carolina Chapel Hill for help with the analysis. We also thank the NYUAD Center for Genomics and Systems Biology for sequencing support and the New York University IT High Performance Computing for supplying the computational resources, services, and staff expertise.

## Supporting information

S1 Fig. ADMIXTURE plot of K=2 to 7 for the 40 *M. polymorpha* individuals.

S2 Fig. Phylogenetic relationship of *M. polymorpha*, *M. rugosa*, and *M. tremuloides*.

S3 Fig. Treemix graph of the Hawaii-Island *M. polymorpha* and its outgroup from m=0 to m=5.

**S4 Fig. Plot of squared correlation coefficient (r^2^) between polymorphic sites.** Decay of linkage disequilibrium was plotted by averaging r^2^ in 100-bp bins.

S5 Fig. Genome-wide F_ST_ calculated in 10-kbp windows between G_H1_ and N.

S6 Fig. Visual representation of the 20 demographic models.

**S7 Fig. dadi model fit for the best-fitting demographic model.** Above row shows the observed and model fit folded site frequency spectrum. Below shows the map and histogram of the residuals.

S8 Fig. Visual representation of the 7 vicariance demographic models.

S9 Fig. Levels of polymorphism (θ_π_) between islands of divergence (zF_ST_ > 4) and the genomic background (zF_ST_ < 4) between G_H1_ and N.

S10 Fig. Plot of the proportion of genic sequences to f_4_ statistics of the topology [G_H1_,N;G_M_,T] in 10-kbp windows.

S1 Table. Information on the sequenced and analyzed individuals of this study.

**S2 Table. dadi parameter estimates for 20 different models using genotype likelihood based 2D-SFS for the G_H1_ and N populations.** nu1,nu2: population 1 and 2 effective population size under models without a population size change; nu1a, nu2a: population 1 and 2 effective population size before a population size change; nu1b, nu2b: population 1 and 2 effective population size after a population size change; m12, m21: migration rate. For symmetric migration models only m12 is shown; m12a, m21a: migration rate of first epoch; m12b, m21b; migration rate of second epoch; T1,T2, T3: divergence times for different epoch periods of the model.

**S3 Table. dadi parameter estimates for 20 different models using genotype call-based 2D-SFS for G_H1_ and N populations.** nu1,nu2: population 1 and 2 effective population size under models without a population size change; nu1a, nu2a: population 1 and 2 effective population size before a population size change; nu1b, nu2b: population 1 and 2 effective population size after a population size change; m12, m21: migration rate. For symmetric migration models only m12 is shown; m12a, m21a: migration rate of first epoch; m12b, m21b; migration rate of second epoch; T1,T2, T3: divergence times for different epoch periods of the model.

**S4 Table. dadi parameter estimate for top 3 best-fitting models after further optimization steps for G_H1_ and N populations.** nu1,nu2: population 1 and 2 effective population size; m12, m21: migration rate. For symmetric migration models only m12 is shown; T1,T2: divergence times for different epoch periods of the model.

**S5 Table. dadi parameter estimates for the 7 vicariance models for G_H1_ and N populations.** nu1,nu2: population 1 and 2 effective population size; m12, m21: migration rate. For symmetric migration models only m12 is shown; T1,T2: divergence times for different epoch periods of the model. s: proportion of ancestral population that lead to population 1, and population 2 was established with 1-s.

**S6 Table. dadi parameter estimates for the top 3 best-fitting vicariance models after further optimization steps for G_H1_ and N populations.** nu1,nu2: population 1 and 2 effective population size; m12, m21: migration rate. For symmetric migration models only m12 is shown; T1,T2: divergence times for different epoch periods of the model. s: proportion of ancestral population that lead to population 1, and population 2 was established with 1-s.

**S7 Table. dadi parameter estimates for 20 different models using genotype call-based 2D-SFS for G_H2_ and N populations.** nu1,nu2: population 1 and 2 effective population size under models without a population size change; nu1a, nu2a: population 1 and 2 effective population size before a population size change; nu1b, nu2b: population 1 and 2 effective population size after a population size change; m12, m21: migration rate. For symmetric migration models only m12 is shown; m12a, m21a: migration rate of first epoch; m12b, m21b; migration rate of second epoch; T1,T2, T3: divergence times for different epoch periods of the model.

**S8 Table. dadi parameter estimate for top 3 best-fitting models after further optimization steps for G_H2_ and N populations.** nu1,nu2: population 1 and 2 effective population size; m12, m21: migration rate. For symmetric migration models only m12 is shown; T1,T2: divergence times for different epoch periods of the model.

S9 Table. Median D_xy_ within and outside the genomic islands of divergence found in G_H1_ and N for various population comparisons.

S10 Table. List of gene names overlapping genomic islands of divergence.

